# Critiquing Protein Family Classification Models Using Sufficient Input Subsets

**DOI:** 10.1101/674119

**Authors:** Brandon Carter, Maxwell Bileschi, Jamie Smith, Theo Sanderson, Drew Bryant, David Belanger, Lucy Colwell

**Author notes:** Correspondence to: Brandon Carter < >.

## Abstract

In many application domains, neural networks are highly accurate and have been deployed at large scale. However, users often do not have good tools for understanding how these models arrive at their predictions. This has hindered adoption in fields such as the life and medical sciences, where researchers require that models base their decisions on underlying biological phenomena rather than peculiarities of the dataset introduced. In response, we propose a set of methods for critiquing deep learning models and demonstrate their application for protein family classification, a task for which high-accuracy models have considerable potential impact. Our methods extend the sufficient input subsets technique, which we use to identify subsets of features (SIS) in each protein sequence that are alone sufficient for classification. Our suite of tools analyzes these subsets to shed light on the decision-making criteria employed by models trained on this task. These tools expose that while deep models may perform classification for biologically-relevant reasons, their behavior varies considerably across choice of network architecture and parameter initialization. While the techniques that we develop are specific to the protein sequence classification task, the approach taken generalizes to a broad set of scientific contexts in which model interpretability is essential.

## 1. Introduction

In recent years, deep neural networks (DNNs) have provided considerable performance improvements for a wide variety of machine learning (ML) prediction problems. However, their adoption has been slower in applied domains such as the life and medical sciences, where practitioners often require that models make decisions for biologically relevant reasons. Without this property, models trained on experimentally collected data cannot be trusted, since they may be fitting to systematic biases introduced during data collection. Therefore, there is demand for tools that help evaluate the consistency between prior knowledge about how predictions should be made and the mechanisms actually employed by a given ML model in practice.

This paper presents a set of analytical tools for validating model behavior that extend the *sufficient input subsets* method (SIS) (Carter et al., 2019). Suppose a model predicts *y* given an input *x*, where *x* is multi-dimensional, such as a sequence of characters. SIS finds a collection of minimal subsets of the components of *x* such that the model makes the same prediction using any of the subsets alone. The sufficient input subsets are small, human-readable compressions of *x* that allow easy-to-visualize analysis. One context where SIS is particularly useful is for interrogating models trained on biological sequence data, where the resulting subsets can be reconciled with external knowledge about the underlying dependence of *y* on *x*.

Our experiments focus on models that predict the functional classification of a protein sequence from its raw amino acid sequence. A protein is synthesized as a linear sequence of amino acids. This sequence typically contains the information required to specify the 3D tertiary protein structure, and moreover, the function of the resulting molecule. The falling cost of high-throughput sequencing means that sequences of novel proteins are quickly accumulating in databases. However, the cost and expertise required to experimentally characterize these proteins lags far behind (Price et al., 2018). This means that the vast majority of protein sequence annotations are obtained using computational models, which have limited coverage. For example, one-third of all protein-coding genes from bacterial genomes currently cannot be annotated with a function using current models (Chang et al., 2015), suggesting that new approaches could have significant impact. Besides categorizing existing proteins, such models also have the potential to guide the discovery of novel proteins that catalyze reactions, are important for biotechnology, or produce new therapeutics.

Hidden Markov models (HMMs) provide state of the art performance (Finn et al., 2011) for this task, but there are suggestions in the literature that DNNs have the potential to provide a new modeling paradigm (Hou et al., 2017; Seo et al., 2018; Dalkiran et al., 2018; Li et al., 2017a; Liu, 2017; Bileschi et al., 2019). In particular, HMMs do not capture interactions between non-local positions in protein sequence, which have recently been shown to be highly predictive of protein tertiary structure (Marks et al., 2011; Xu, 2018; R.Evans, 2018) and function (Bitbol et al., 2016; Gueudré et al., 2016; Riesselman et al., 2018).

In the literature, deep models have been shown to achieve extremely high accuracy on held-out data (Bileschi et al., 2019), to accurately label remote homologs (Li et al., 2017a), and in a handful of cases their predictions have been experimentally validated (Liu, 2017). Unfortunately, we are currently unable to characterize their behavior, leading to lack of trust among practitioners (Angermueller et al., 2016; Montavon et al., 2018; McCloskey et al., 2018). This motivates our development of tools to interrogate these complex black-box functions, since traditional metrics like held-out test accuracy do not provide assurance that the models can be used as effective surrogates for laboratory experiments.

In response, we present six novel extensions of SIS that allow us to inspect and critique the decision-making of protein sequence classification models. Our contributions include:

- **SIS-3D**: We show the sufficient input subset of a particular input protein domain rendered in 3 dimensions, as a subset of a folded protein (Sections 4.1, 5.1).
- **SIS logo**: We aggregate sufficient input subsets across many classifications, generating a *SIS logo* visualization, which allows us to globally inspect model decision-making (Sections 4.1, 5.2).
- **Explaining misclassifications**: We provide an interpretable, numerical measure for understanding why a model made a particular classification, as opposed to a different choice (Sections 4.1, 5.3).
- **Feature compression**: We compare different models’ abilities to rationalize decisions across a large number of inputs, contrasting SIS size needed for a particular confidence level in the decisions (Sections 4.1, 5.4).
- **SIS location**: We compare different models’ considerations of amino acid placement within sequences, and show that different model architectures favor features at different sequence regions (Sections 4.1, 5.5).
- **Model inconsistency**: We show that identical models, when retrained, have different rationales for their decision-making and quantify this instability (Sections 4.1, 5.6).

## 2. Background and Related Work

In this section, we describe our approach for model interpretation and review existing literature on deep neural networks applied to protein sequence data.

### 2.1 Methods for Interpreting ML Predictions

A wide range of methods have been developed to help practitioners interpret the often complex behavior of machine learning models. Common approaches to interpret model behavior include use of architectures that produce human-interpretable prediction models, such as attention mechanisms (Sha & Wang, 2017) or the generator-encoder approach of Lei et al. (2016). However, in practice, these techniques are not applicable in situations where achieving state of the art accuracy (Caruana et al., 2015) requires a specific model parameterization or because the model is only accessible via a black-box API (for example, a model which can be queried through a public API and whose parameters and data are hidden) (Tramèr et al., 2016). Other model-agnostic interpretability approaches produce attribution scores that quantify the importance of each feature in determining the output of a function *f* on an input *x*. Such methods include LIME (Ribeiro et al., 2016), which fits a local linear model to *f(x)*, saliency maps based on gradients of *f* (Baehrens et al., 2010; Simonyan et al., 2013), Layer-wise Relevance Propagation (Bach et al., 2015), DeepLIFT (Shrikumar et al., 2017), integrated gradients (Sundararajan et al., 2017), and input-signal-based techniques using gradients of *f* (Zeiler & Fergus, 2014; Springenberg et al., 2014; Kindermans et al., 2017b). However, such gradient-based methods have been shown to be unreliable, depending not only on *f*, but also on its architectural implementation and scaling of inputs (Kindermans et al., 2017a;b).

Another recent approach for interpreting models is a method based on *sufficient input subsets* (SIS) (Carter et al., 2019). SIS is a local explanation framework that produces rationales (known as sufficient input subsets) for a model which maps inputs *x ϵ χ* via a function *f* : *χ* → ℝ. Each sufficient input subset consists of a minimal input pattern present in *x* that alone suffices for *f* to produce the same decision, even with information about the rest of the values in *x* missing. SIS presumes that the decision is based on *f(x)* exceeding some pre-specified threshold *τ* ∈ ℝ and identifies a complete collection (*SIS-collection*) of disjoint subsets *S*, each satisfying *f(x*_*S*_ ) ≥ *τ*. As in Carter et al. (2019), we use the notation *x*_*S*_ to represent a modified version of *x* in which all information about the features outside the subset *S* has been masked, while features in *S* retain their original values. Briefly, the SIS method identifies such collections by iteratively applying backward selection to each example *x*. At each step, the algorithm identifies a minimal subset of features *S*_*k*_ that meets the decision criterion 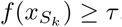, then masks the features in *S*_*k*_ from *x* and repeats the process to identify additional such subsets. Because each SIS would on its own lead the model to reach the same decision, the decision made for an input *x* can be understood through its SIS-collection.

Unlike many other methods, SIS can be applied to interpret any black-box model, is completely faithful to the underlying function *f*, does not require differentiability or gradient information from *f*, does not use any auxiliary explanation model, and can be easily applied and understood by non-experts. Furthermore, unlike many attribution methods that consider each input variable in isolation, SIS operates in a manner that allows interactions between variables that are important for model decision-making to be captured in the sufficient input subset. This is particularly relevant for the domain of protein sequence data, where non-local interactions between sequence positions are known to be important for determining protein structure and function. In this paper, we use and extend the SIS method to interpret protein family classification models.

### 2.2 Neural Networks for Proteins

Recent work has demonstrated that deep learning models have significant potential for the accurate classification of protein sequences. For example, DeepFam (Seo et al., 2018) uses a single convolutional layer and a single fully connected layer to classify sequences from 2892 COG families each containing more than 100 sequences, showing comparable performance to profile HMMs (pHMMs). Similarly, DeepSF is a 30-layer convolutional neural network (CNN) that classifies sequences from 1195 fold classes taken from SCOP, reporting better fold recognition abilities than the state of the art tool HHsearch, though the deep models did not perform so well on finer-grained superfamily and family levels (Hou et al., 2017; Söding, 2004; Andreeva et al., 2004). A deep CNN has also been trained to annotate sequences from SwissProt with hundreds of class labels including some multi-label examples, where the minimum training class size considered is 150 (Szalkai & Grolmusz, 2018; Consortium, 2016). The performance of recurrent neural networks (RNNs) for classifying sequences from four functional classes has also been experimentally validated (Liu, 2017). Similarly, a model that combines a CNN with an RNN was used to predict enzyme classifications (Li et al., 2017b). Finally, Bileschi et al. (2019) demonstrate that a dilated convolutional neural network can annotate protein function from amino acid sequence as accurately as BLASTp (Altschul et al., 1997) and pHMMs in a classification task on Pfam (El-Gebali et al., 2018) sequences. This previous work demonstrates the potential for deep models to accurately predict the function of sequences that cannot be annotated using existing HMMs.

## 3. Neural Network Architectures and Data

In our experiments, we train and explore three neural network architectures on the protein family classification task: a deep convolutional neural network (*deep CNN*); a simpler, shallower convolutional neural network (*shallow CNN*); and a recurrent neural network (*RNN*). We use the same model architectures and training as Bileschi et al. (2019), adopting the ProtCNN, 1-ResNet Block CNN, and RNN architectures, respectively. In this section, we provide a summary of the data and models employed in our experiments (details in Supplement S1).

### TRAINING DATA

We work with pre-cut sequence domain data from the popular Pfam v32.0 seed database (El-Gebali et al., 2018). A domain is a functional unit found within a protein sequence that is conserved across evolutionary time (Buljan & Bateman, 2009); these domains are akin to the pre-segmented images in ImageNet (Deng et al., 2009), as opposed to natural images that may contain many object classes. We use raw amino acid sequences as the only input features, with no multiple sequence alignment step. Domain sequences can be as short as 25 amino acids, or as long as 1000, but most have length 100-250. Each domain sequence is labelled as belonging to just one of the 17929 families in Pfam v32.0. Each example is a sequence of characters, each of which comes from a vocabulary of the 20 common amino acids. We use the same data as in Bileschi et al. (2019), which is available online.^1^ Pfam seed sequence sets with 10 or more members are randomly split into disjoint train (80%), dev (10%), and test (10%) sets. Those 4858 families that have fewer than 10 seed sequences are only present in the train set. The total number of training examples is 1086741, and the number of dev and test examples is 126171.

### DILATED RESIDUAL CNNS

Both the shallow and deep CNNs are residual networks. In a residual network (He et al., 2016), each layer is an additive shift of the previous layer, with layer *i* having *f*_*i*_ = *f*_*i-1*_ *+ g*_*i*_*(f*_*i-1*_). Here, each *f*_*i*_ is an *L* × *F* array and *g*_*i*_(·) is a one-layer convolutional network. The models are easier to train in practice than traditional architectures because they are less susceptible to vanishing gradients (He et al., 2016). The deep CNN also employs dilated convolutions, which help capture long-range interactions in the inputs (Yu & Koltun, 2016). The kernels have “holes” in them, so that the number of trainable parameters does not scale with the convolution’s receptive field size. We use a dilation rate of 2 in each residual block, such that the model’s overall receptive field size is exponential in its depth.

### BIDIRECTIONAL LSTMS

The RNN is a one-layer bidirectional RNN with LSTM cells (Hochreiter & Schmidhuber, 1997). In a bidirectional RNN, we apply two unidirectional RNNs to the sequence of *L* amino acids, one running forwards along the sequence and one in reverse, yielding two arrays of shape *L* × *F*. These arrays are concatenated to produce an output of shape *L* × *2F.*

### POOLING AND PREDICTION

Each of the three model architectures produces a matrix of shape *L × F* for a sequence of length *L* and some *F* (the number of features). This output is mapped to a vector of length *F* by pooling along the length of the sequence. For both CNNs, we use max pooling, which takes the maximum activation along the *L* axis of the array. For the RNN, we use mean pooling, which takes the mean activation along the *L* axis. A linear layer and softmax is then applied, yielding a vector of predicted probabilities for each class (protein family). All models are trained on the Pfam seed train set using Adam (Kingma & Ba, 2014). Hyperparameters are tuned using random sampling (CNNs) or Bayesian optimization using Gaussian process regression (Golovin et al., 2017) (RNN) to maximize accuracy on validation data (Table S1).

## 4. SIS for Protein Sequence Classification

For a given protein sequence *x* and trained model *f*, we apply the SIS procedure to produce a SIS-collection of subsets *S* of sequence positions in *x*. To mask a position in a protein sequence as required by SIS, we replace the amino acid with the X character (a standard representation for “any amino acid”). Our models consume one-hot representations for the 20 common amino acids, and thus we represent the mask X as a vector consisting of the value 1/20. This approach is similar to that in Carter et al. (2019) for masking DNA sequences.

SIS requires a scalar-valued output of the model, whereas our models output a vector of probabilities for each class. In most contexts below, we define *f* to return the scalar output probability corresponding to the predicted (most probable) class. Across all of our experiments we set *τ* ≥ = 0.9 as the probability threshold for deciding that a sequence belongs to a family, i.e. we require *f*(*x*) 0.9. SIS then returns a complete collection of sufficient input subsets, each of which satisfies *f(x*_*S*_ ) ≥ 0.9. Note that here *x*_*S*_ is a masked version of the initial sequence *x* in which only sequence positions in *S* are included, and all other elements are masked. We verified that the distribution over classes for a fully-masked sequence *x*_all mask_ (a sequence consisting of only X characters) was nearly uniform and that max_*i*_[*f*_*i*_(*x*_all mask_)] ≪ 0.9, where *f*_*i*_*(x)* is the predicted probability of class *i*.

### 4.1 Methodological Contributions

Here, we introduce six extensions of the SIS method, which we use to interrogate the behavior of our protein classification models.

#### SIS-3D

The sufficient input subsets returned by SIS can be understood as justifications for a classifier’s decisions on individual sequences. Viewing these subsets in sequence space is of limited utility, because the sequence position of an amino acid generally has little relationship to the functional role that it plays. In contrast, the location of the SIS positions in the three-dimensional (3D) structure of the folded protein contains much more information about their involvement in protein function (Berezin et al., 2004; Marks et al., 2011). For example, amino acids that line the substrate binding pocket of a protein often play a central role in substrate recognition, while those found at a protein binding interface contribute significantly to protein interaction specificity. Therefore, interpretability is significantly improved by locating these amino acids within a 3D rendering of the folded protein structure. We use PyMOL (Schrödinger, LLC, 2015) together with coordinates from the solved protein structure 1a3n in the protein data bank (Burley et al., 2017; Tame & Vallone, 2000) to render this visualization (see Figure 1).

**Figure 1.**
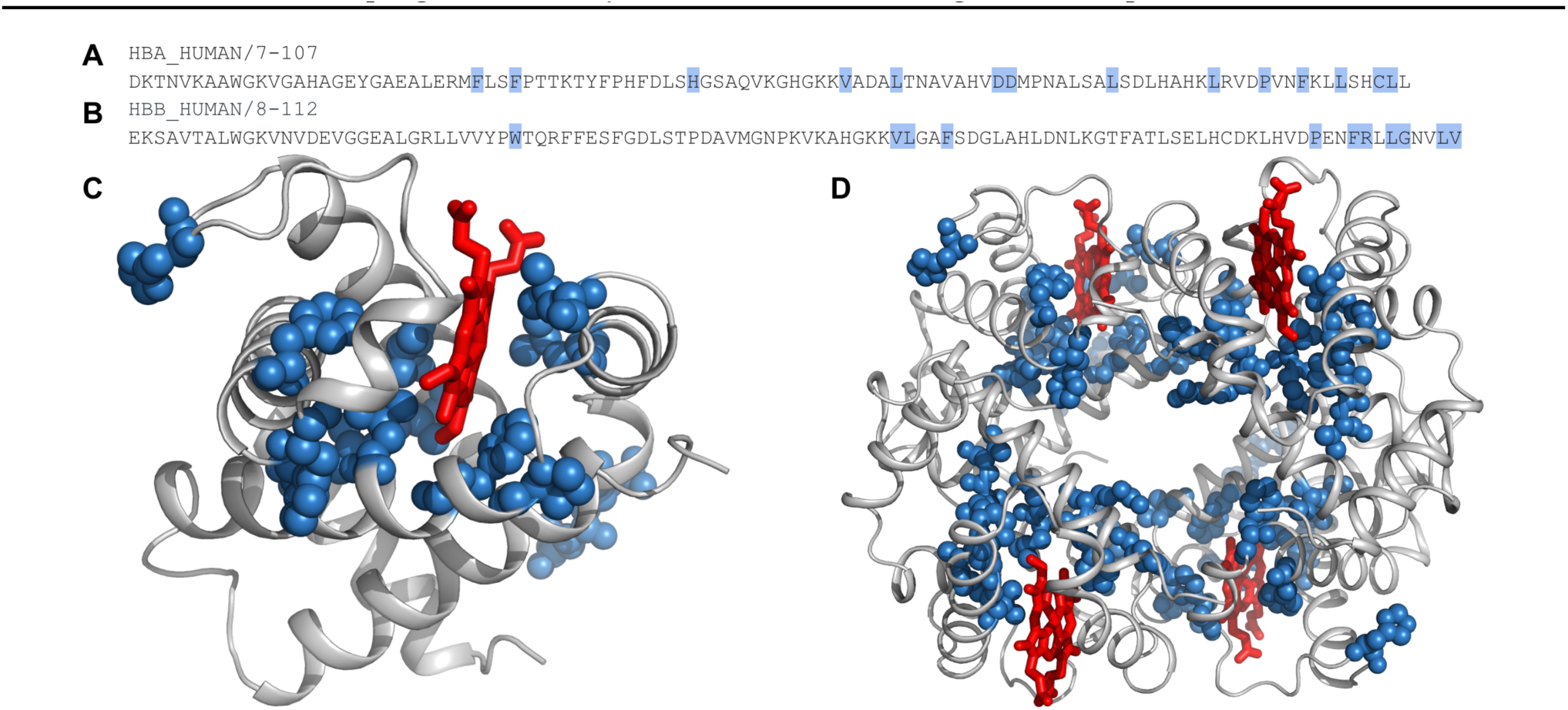
(a) Domain sequence HBA HUMAN/7-107, SIS from a deep CNN is shaded. (b) Domain HBB HUMAN/8-112, SIS from a deep CNN is shaded. (c) The HBA subunit of human hemoglobin, with the SIS from (a) rendered as spheres. Note the proximity to the heme group (rendered in red sticks). (d) Human hemoglobin, including two copies each of the HBA and HBB subunits. SIS amino acids from (a) and (b) shown as spheres, heme substrate rendered using red sticks.

#### SIS LOGO

We introduce an approach for visualizing the set of SIS collections found across all sequences that belong to a particular family. The resulting visualization provides an immediate visual summary of all SIS associated with the family, and moreover the level of variation among SIS within that family. Our approach builds on the concept of sequence alignment, which is central to current protein sequence analysis methods. Specifically, *multiple sequence alignments* (MSAs) are available for all the families in Pfam v32.0 (El-Gebali et al., 2018). This allows us to map each SIS within a family to the same reference alignment, and compute a histogram across the set of SIS-collections for a family. The histogram indicates the frequency with which each amino acid occurs at each aligned sequence position.

An MSA can be represented as a *sequence logo* (Schuster-Boöckler et al., 2004) (example shown in Figure 2a) in which each possible amino acid at each position is represented by its frequency, which is then transformed to the relative entropy compared to the background frequency of amino acid usage, measured in bits. High entropy positions represent conserved protein regions that are generally important for protein tertiary structure and function (Ashkenazy et al., 2016). To produce our *SIS logo*, we align the sequences to the reference family MSA described above, mask the non-SIS amino acids in each sequence with X, and visualize the resulting SIS logo using the Skylign tool (with the “Convert to profile HMM - keep all columns” option) (Wheeler et al., 2014). The SIS logo allows us to confirm concordance with the family HMM logo, indicating that the model learns discriminative features that are conserved across the family (see Section 5.2 for results).

**Figure 2.**
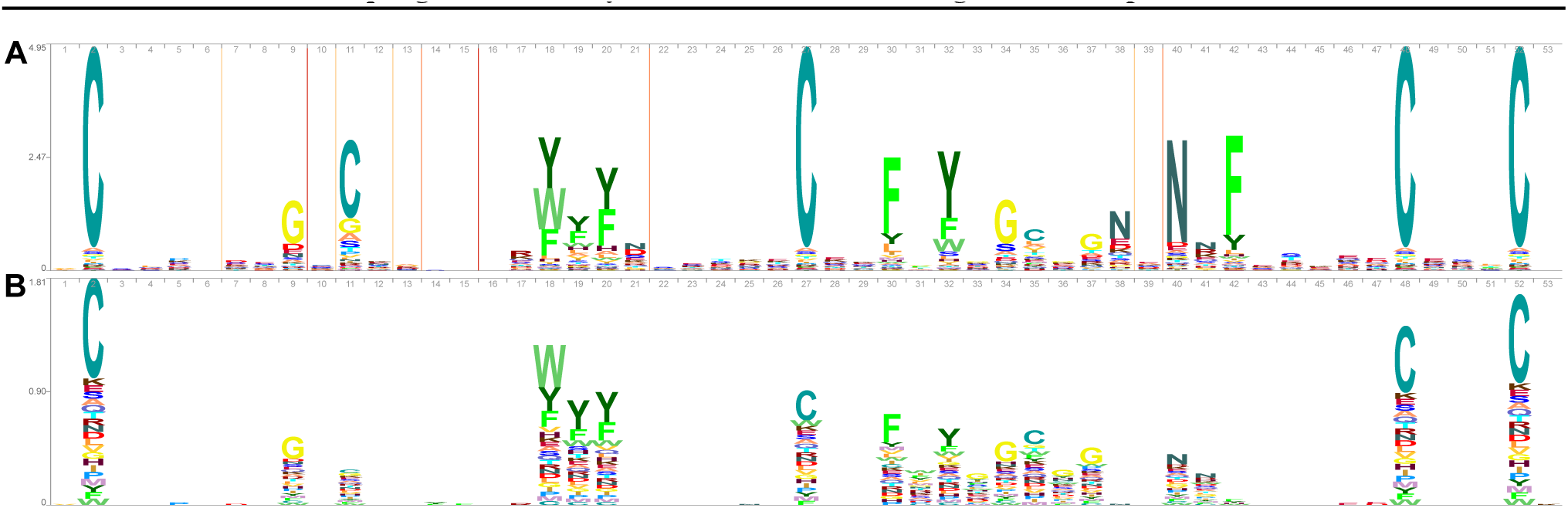
(a) Sequence logo for the Kunitz domain family (PF00014) and (b) SIS logo (see Section 4.1) showing SIS identified for sequences in this family with the deep CNN.

#### EXPLAINING MISCLASSIFICATIONS

We compare the SIS-collection of a misclassified sequence to SIS-collections of sequences from both its predicted and true families. For each of the true and predicted families, we report (as in Figure 3a) the distribution of edit distances from the misclassified sequence’s SIS to all members of that family and the closest SIS from any member of that family to the misclassified sequence’s SIS.

**Figure 3.**
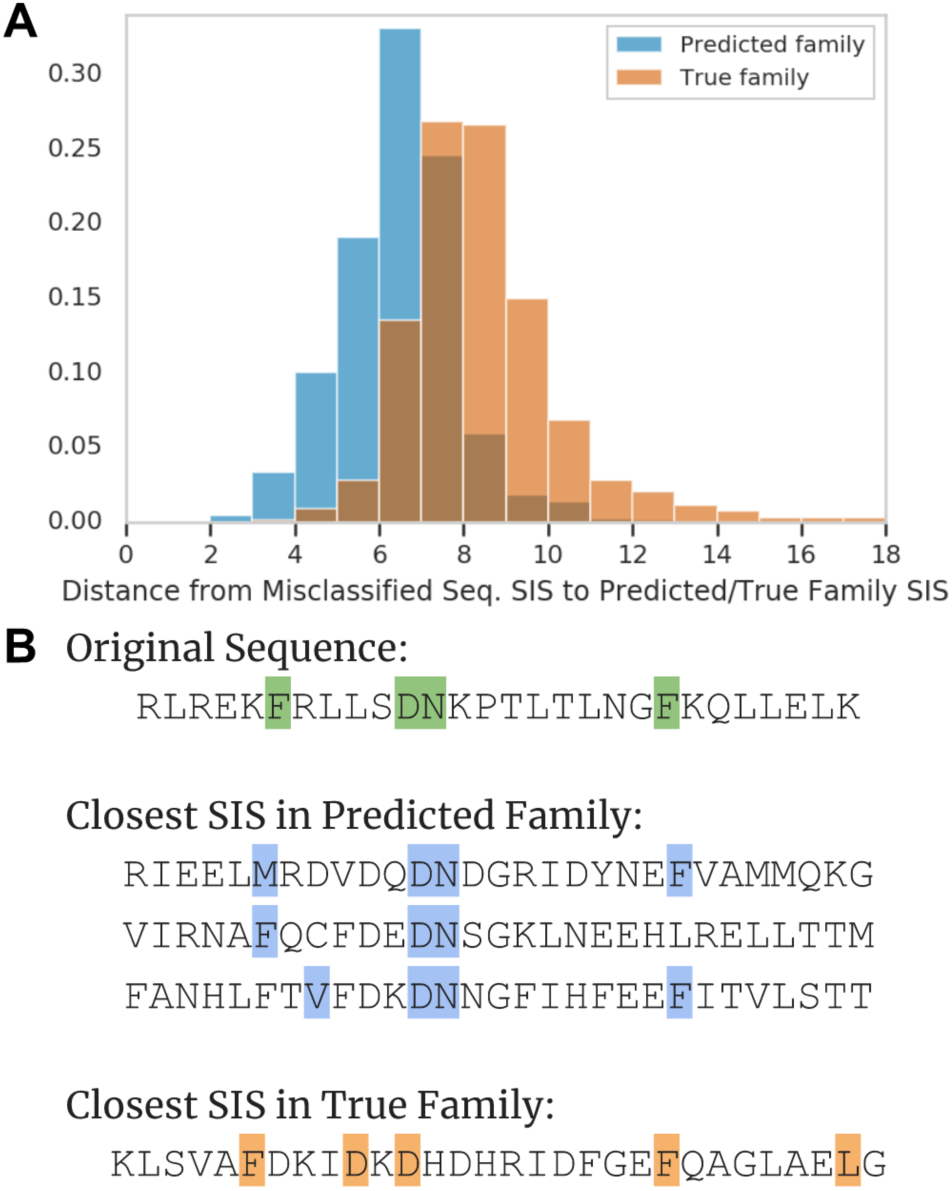
(a) Histogram of edit distances from SIS for a sequence misclassified by the deep CNN to SIS for sequences from the true and predicted families. (b) SIS shown for misclassified sequence as compared to closest SIS from predicted family (distance of 2) and true family (distance of 3). SIS characters are shaded.

Distance is computed as edit (Levenshtein) distance between sequences in which the non-SIS amino acids of each sequence are replaced with X. In contrast to Carter et al. (2019), which only provides a qualitative measure of why a sequence is misclassified, our approach provides a numerical measure of the incorrect decision (see Section 5.3).

#### FEATURE COMPRESSION

Here, we compare models through comparing the succinctness of the rationales for their decisions. For each protein sequence *x* and SIS *S* in its SIS-collection, we compute the fraction of *x* comprising *S* as |*S* | */* |*x* |, where |*S* | is the number of positions in *S* and |*x* | is the length of *x*. Then, for each (SIS-collection, sequence, architecture, replicate) pairing, we compute the fraction |*S* | */* |*x* | of the sequence positions in each SIS of the collection. See Section 4.2 for discussion of model replicates and Section 5.4 for results.

#### SIS LOCATION

We examine whether there are differences in which sequence positions, or locations, are used by the different model architectures to classify sequences. We observe which positions are captured in each SIS, and then normalize this positional information to relative position in the sequence. As above, we repeat this experiment across SIS from retrained models for each architecture (see Section 4.2). See Section 5.5 for results.

#### MODEL INCONSISTENCY

One point of potential concern for practitioners is the stability of models: does the same architecture retrained on the same data (with different random initialization) make the same decisions? Are those decisions made for the same reasons? In order to address these questions, we compare two replicates of the same architecture and use each to make predictions on the SIS from the other model, as in Carter et al. (2019). Here, we also compare SIS from each of the models on the same examples to evaluate whether the models are consistent in rationalizing predictions on specific instances (see Section 5.6).

### 4.2 Engineering and Implementation Details

Our implementation of SIS is available on GitHub.^2^

#### MODEL REPLICATES

To reduce the effect of noise, we retrain each of our architectures 10 times with different random parameter initializations and input sampling during training. This allows us to estimate variance in our quantitative analyses and be more confident in concluding that the results are inherent to an architecture, not its initialization.

#### COMPUTING 4 BILLION MODEL PREDICTIONS

One of the principal challenges in implementing the methods described in Section 4.1 is the computational cost of running SIS across a large number of sequences (∼ 10k), model architectures (3), and replicates of each architecture (10). The average sequence length in our evaluation set is about 144 characters. For each sequence, the lower bound on the number of inferences (forward propagation in the neural networks) needed to compute its SIS-collection is 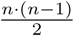, where *n* is the sequence length. This gives us 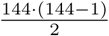, approximately 4 billion inferences. As with any experimentation, we had to be able to iterate and improve our methodology, leading to running this 4-billion-inference experiment many times.

This computational challenge motivates a significant engineering effort to run the SIS procedure in a distributed environment. We use the Apache Beam Python SDK, a MapReduce (Dean & Ghemawat, 2008) implementation based on Akidau et al. (2015). Specifically, we leverage only the Map and Shuffle primitives to distribute tasks among workers. Since the computation of each individual sequence’s SIS is an embarrassingly parallel operation, we group each input example (a sequence) into its own group, and then the shuffler distributes these groups equally to the maximum number available reducers. This allows us to achieve maximum parallelization when running SIS. Because CPUs are much more widely available and are cheaper than GPUs, we run inference on CPUs. As with most MapReduce applications, we face a long tail of slow computations. We found that, as SIS is runs in *𝒪(n*^*2*^) time for a sequence in length *n*, long sequences disproportionately affect the running time of our Beam jobs.

#### ONE PERCENT SAMPLE

In order to limit the number of inferences we had to perform, instead of computing a SIS for each member of Pfam seed (El-Gebali et al., 2018), we uniformly sample 1%, and use this set for some of our analysis. For the reasons described above, we exclude sequences longer than 500 amino acids (∼ 2% of sequences) from this sample. The resulting sample contains 13188 sequences from 5933 unique protein families. Note that models are trained on the entire train set.

## 5. Results

In this section, we evaluate DNNs trained to classify protein sequences (see Section 3). SIS allows us to characterize model behavior both locally (on individual examples) as well as globally (through family-level SIS logo visualizations). Our methods enable us to critique and compare their behavior.

### 5.1 SIS-3D

We first focus on the deep CNN model (Section 3) and show how sufficient input subsets can be understood as justifications for the CNN’s decisions on individual sequences. The shaded amino acids in Figures 1a, b display sufficient input subsets for the human hemoglobin *α* and *β* (3 domain sequences, which were correctly classified by the CNN as members of the globin family (PF00042). We note that these subsets contain roughly 10%–15% of the sequence positions, which are scattered throughout the domain sequence. However, distant positions in the sequence may become close when the protein folds into a 3D structure. Figure 1c shows the amino acids in the SIS for human hemoglobin using all-atom sphere representation (blue), superimposed on a representation of the protein backbone (grey). The core function of hemoglobin is to transport oxygen, which binds to the iron atom contained in each associated heme group (red). The heme group binds to a pocket in the 3D structure of the hemoglobin α molecule. Figures 1c, d show that the amino acids contained in the sufficient input subsets for both human hemoglobin *α* and *β* (3 are clustered around the heme binding sites in the 3D structure. The heme binding pocket is highly characteristic of members of the globin domain family, and the fact that the sufficient input subset overlaps with this functionally important feature of globin tertiary structure suggests that the trained model has learned non-trivial information about the globin domain.

### 5.2 SIS Logos

We compute SIS-collections for a large number of sequences from the Pfam v32.0 full dataset (El-Gebali et al., 2018) predicted to belong to the Kunitz domain family (PF00014) with high confidence (probability ≥ 0.9) by the deep CNN. We display the corresponding SIS logo in Figure 2b. We observe a high degree of agreement between the prominent sequence positions in the SIS logo and the highly-conserved positions in the multiple sequence alignment for this family (Figure 2a).

Different regions of protein domains are subject to different constraints. Some sections of a protein may not play a major functional role, and the exact amino acids at these positions may have little effect on protein function. However, a single change to a key amino acid at, for instance, a protein’s active site might fundamentally impair the protein’s ability to perform its role. Traditionally, multiple sequence alignments are used to identify such conserved sequence positions which are likely to be important for protein function (Ashkenazy et al., 2016). Figure 2 suggests that SIS are highly enriched for evolutionarily conserved sequence positions. Note, however, that not all conserved indices appear in the SIS logo. This is because the model learns *discriminative* features in order to distinguish between families. Some sequence positions may be highly conserved, but occur in similar contexts in many families.

### 5.3 Explaining Misclassifications

Our approach also enables us to gain insight into why a model makes particular misclassifications, as seen in Figure 3. The deep CNN misclassifies sequence Q22DM3 TETTS/41-68, a member of family EF-hand_6, as belonging to family EF-hand 1, another helix-loop-helix structure family. We analyze this particular sequence because its predicted probability is nearly 1.0, and the model has few misclassifications from EF-hand 6 to EF-hand 1, and vice versa. Our analysis shows that the SIS for Q22DM3 TETTS/41-68 is closer to SIS of sequences in the predicted family (2 edits) than those from the true family (3 edits). Moreover, Figure 3 shows that the SIS for this misclassified sequence is closer on average to SIS from the predicted family than SIS from the true family, giving further insight into why the sequence was misclassified.

### 5.4 Feature Compression

We next compare our DNN models by evaluating how succinctly they rationalize their decisions. We compute SIS-collections across our 1% sample of Pfam v32.0 seed (El-Gebali et al., 2018) for all replicates of the three DNN architectures (see Section 4.2 for details). Figure 4 shows the distribution of the fraction of the input sequence contained in the SIS (see Section 4.1) for each of our three DNN architectures. This result suggests that the RNN uses significantly fewer sequence positions than the CNN-based architectures to make classifications with the same confidence. Inversely, the deep CNN requires a significantly larger fraction of the sequence to classify proteins than does the shallow CNN. Another way to interpret this result is that the RNN can more succinctly rationalize its decisions than the two CNNs. Whether one desires that a model *should* rationalize its decisions more succinctly or more verbosely likely depends on the precision-recall considerations of a particular application. Our approach provides a method to compare models in this way.

**Figure 4.**
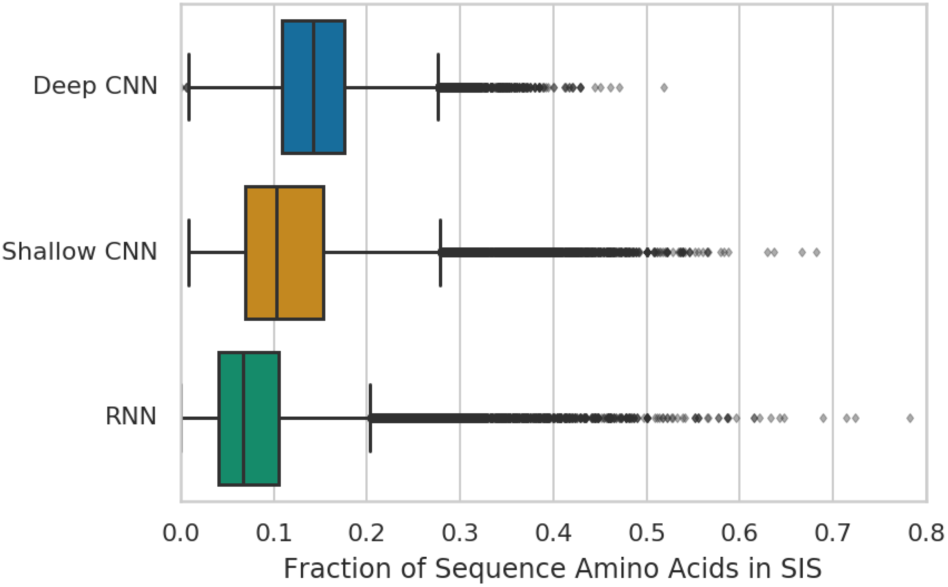
Boxplot indicating fraction of protein sequence positions in each SIS. Distributions are over the union of all SIS from all 10 replicates of each architecture.

### 5.5 SIS Location

We apply our SIS logo methodology to examine whether, in general, different model architectures tend to use sequence positions from different locations to conduct classification. Specifically, we visualize SIS logos using sequences from the Pfam v32.0 full dataset (El-Gebali et al., 2018) classified as lactate dehydrogenase (PF00056) by the deep CNN, shallow CNN, and RNN models (Figures 5b, c, d, respectively). We compare these SIS logos to the family HMM logo (Figure 5a). We observe that while there is significant intersection in the sequence positions comprising sufficient input subsets across these models, positions that are important for RNN classification are found almost exclusively at the start and end of the sequence, whereas the deep CNN often also requires positions in the central region of the domain.

**Figure 5.**
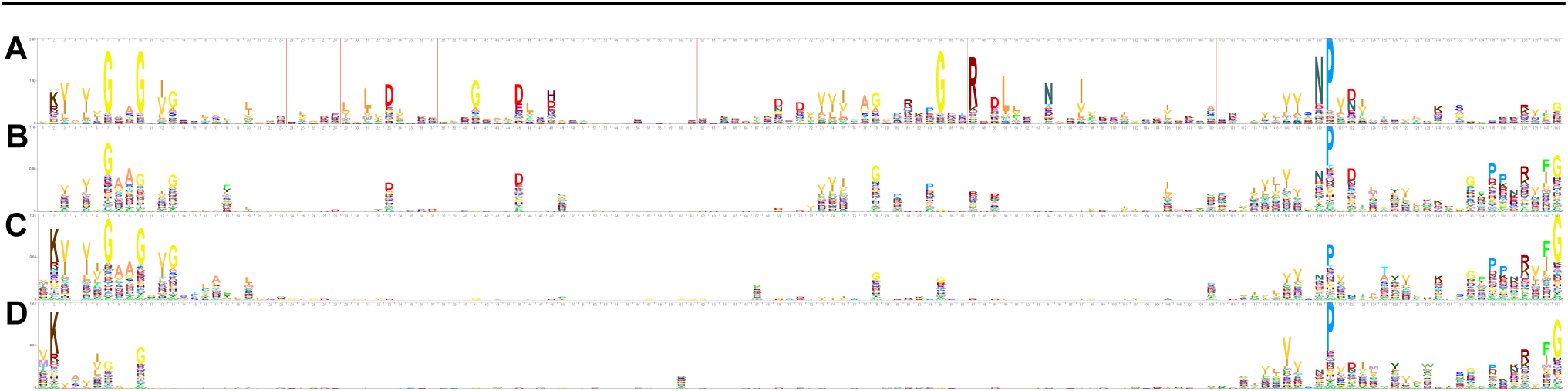
(a) Sequence logo for the lactate dehydrogenase family (PF00056) from Pfam. SIS logos (see Section 4.1) for all SIS identified for this family from (b) deep CNN, (c) shallow CNN, and (d) RNN models.

To assess whether this is specific to this protein family, or a more general property of our trained model architectures, we look at SIS-collections from our random sample of the Pfam seed dataset (see Section 4.2). We observe which positions are captured in each SIS, and then normalize this positional information to a relative position in the sequence.

Aggregating these positional data in Figure 6 reveals that the pattern observed for PF00056 is common across the entire dataset, and reflects a difference in preference for these three different DNNs. To determine whether this is a stochastic feature of training or driven by the model architecture itself, we examine the 10 replicate models trained for each architecture with different random initialization (see Section 4.2). The positional biases of different replicates of each architecture are remarkably consistent (note error bars in Figure 6), with all architectures biased to features at the ends of the sequence, but with this effect stronger in the shallow CNN than the deep CNN, and stronger still in the RNN. This may reflect the difficulties that recurrent models can have in learning long range dependencies. It is also likely that the hand-segmented nature of our training data means that the edges of sequences carry more information, imparted by the curators’ segmentation decisions.

**Figure 6.**
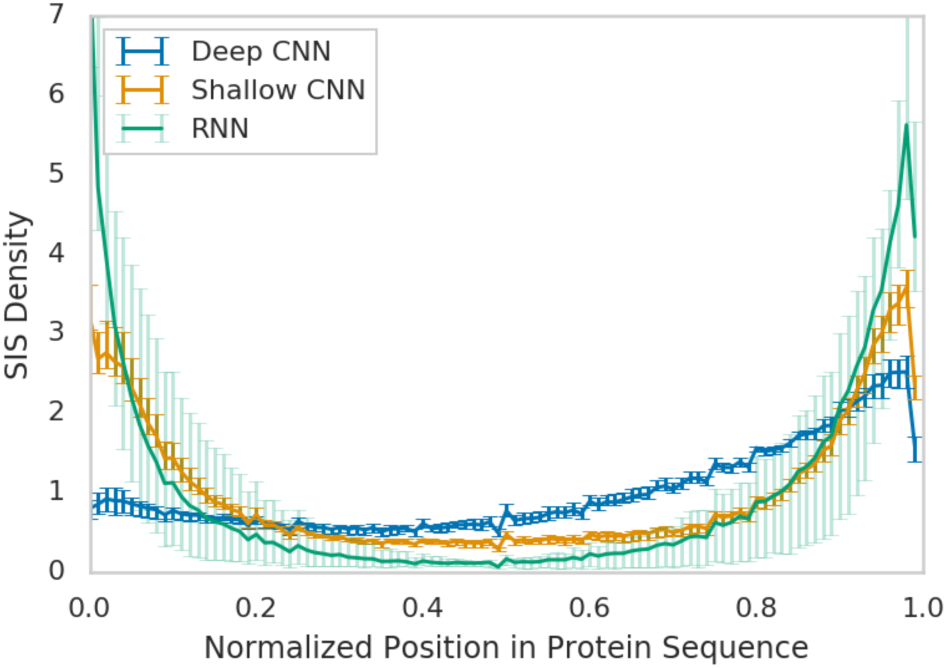
Density of SIS positions throughout protein sequences. For each of the three architectures, we plot the mean density at each position taken over 10 replicate models trained for each architecture. Error bars indicate the minimum and maximum values over the replicates.

### 5.6 Model Inconsistency

As discussed in Section 4.1, one concern for practitioners (e.g. biologists) when considering deep learning models is stability when retrained on the same data. In order to address this question, we train 10 replicate models for each architecture (see Section 4.2). As seen in Table 1, the small variance in accuracy we observe across the replicates of each architecture trained with different random initialization may suggest that the replicate models behave similarly. This is satisfying, given that each DNN training run performs non-convex optimization from a different random initialization. However, while the models seem to be stable in terms of their outputs, they may still have differences in terms of the specific decision boundaries that they learn from the training data, and hence, may classify sequences for different reasons. We probe this question by evaluating whether the rationales employed for arriving at these predictions are consistent across the replicates.

**Table 1.** Final test set accuracies of our trained models. Each value indicates mean over the 10 replicate models per architecture (see Section 4.2) *±* 2 σ

| ARCHITECTURE | ACCURACY (%) |
| --- | --- |
| DEEP CNN | 99.3 ± 0.03 |
| SHALLOW CNN | 98.9 ± 0.08 |
| RNN | 97.7 ± 1.54 |

We choose two deep CNN models (selected at random from the 10 replicates) and use each to make predictions given the SIS from the other model (using all SIS for sequences in our random sample of the Pfam seed dataset, Section 4.2). Figure 7a shows the distributions of these predictions for the two models and suggests that the models are in fact classifying sequences for different reasons. Figure 7b shows an example of a single protein sequence, F1QK57 DANRE/2132-2166 (chosen at random), for which SIS identifies a SIS-collection with one sufficient input subset for each of the models. The shaded characters indicate the SIS for this sequence for each model, such that each model classifies the sequence with confidence 0.9 using just the shaded amino acids. As seen, there is little overlap between the shaded subsets, suggesting the two models (with identical architectures) are learning different decision boundaries to classify sequences. In response to this instability, our analysis in Section 5.4 and 5.5 averages over multiple model replicas.

**Figure 7.**
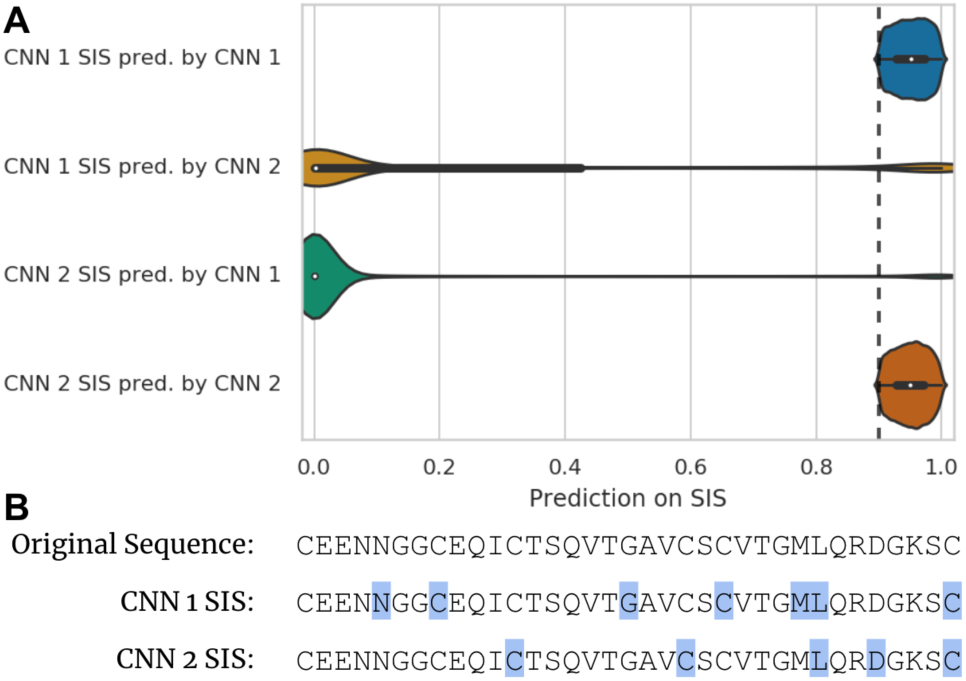
(a) Violin plot showing the predictions by one deep CNN on the SIS extracted from another deep CNN. Dashed line indicates the SIS threshold (τ = 0.9). (b) SIS computed for each of the two CNNs on a randomly chosen sequence (SIS positions are shaded).

## 6. Discussion

We contribute a diverse set of analytical tools that employ SIS to critique the behavior of deep models for protein domain sequence classification. These allow us to understand the fidelity of the models to the underlying biology and to interrogate their stability and reliability. Many of these techniques are very general, and could be applied more broadly to other machine learning models across a broad range of application domains.

We find evidence that our trained models make predictions for biologically-relevant reasons (Sections 5.1 and 5.2) and that model misclassifications can be interpreted (Section 5.3). However, we find that a sequence can be accurately classified using a small subset of its positions (Section 5.4) and the subsets chosen vary across different instantiations of the same model (Section 5.6). Consequently, a model can classify a sequence accurately while e.g. ignoring all but its beginning and end (Section 5.5), suggesting that the set of features that dictates family membership is quite highly redundant.

Depending on their specific use-cases for such a model, researchers may use the above analyses to conclude that a model can or cannot be trusted. In certain contexts, the researcher may simply require that the model achieves high accuracy and makes predictions for sensible reasons. Our models seem to achieve this. On the other hand, researchers sometimes use predictive models for additional higher-level tasks, e.g. to perform *in-silico* screening to estimate the functional impact of mutations. Here, the model is treated as a surrogate to a laboratory experiment and is evaluated on both the original sequence occurring in the wild and on mutations of the sequence. The experiment will be unreliable if the models’ rationales are not scientifically sound, which may be the case if for example predictions are unstable across different initializations. Furthermore, the model may underestimate the impact of mutations in the middle of the sequence. Our techniques can be used to uncover such insights.

## Acknowledgements

We thank Zachary Nado and D. Sculley for helpful comments on this paper.

## Footnotes

1 https://www.kaggle.com/googleai/pfam-seed-random-split

2 https://github.com/google-research/google-research/tree/master/sufficient_input_subsets

## References

Akidau, T., Bradshaw, R., Chambers, C., Chernyak, S., Fernández-Moctezuma, R. J., Lax, R., McVeety, S., Mills, D., Perry, F., Schmidt, E., et al. The dataflow model: a practical approach to balancing correctness, latency, and cost in massive-scale, unbounded, out-of-order data processing. Proceedings of the VLDB Endowment, 8(12):1792–1803, 2015.

Altschul, S. F., Madden, T. L., Schäffer, A. A., Zhang, J., Zhang, Z., Miller, W., and Lipman, D. J. Gapped blast and psi-blast: a new generation of protein database search programs. Nucleic acids research, 25(17):3389–3402, 1997.

Andreeva, A., Howorth, D., Brenner, S. E., Hubbard, T. J., Chothia, C., and Murzin, A. G. Scop database in 2004: refinements integrate structure and sequence family data. Nucleic acids research, 32(suppl 1):D226–D229, 2004.

Angermueller, C., Pärnamaa, T., Parts, L., and Stegle, O. Deep learning for computational biology. Molecular systems biology, 12(7):878, 2016.

Ashkenazy, H., Abadi, S., Martz, E., Chay, O., Mayrose, I., Pupko, T., and Ben-Tal, N. Consurf 2016: an improved methodology to estimate and visualize evolutionary conservation in macromolecules. Nucleic acids research, 44(W1):W344–W350, 2016.

Bach, S., Binder, A., Montavon, G., Klauschen, F., Müller, K.-R., and Samek, W. On pixel-wise explanations for non-linear classifier decisions by layer-wise relevance propagation. PloS one, 10(7):e0130140, 2015.

Baehrens, D., Schroeter, T., Harmeling, S., Kawanabe, M., Hansen, K., and MÃžller, K.-R. How to explain individual classification decisions. Journal of Machine Learning Research, 11(Jun): 1803–1831, 2010.

Berezin, C., Glaser, F., Rosenberg, J., Paz, I., Pupko, T., Fariselli, P., Casadio, R., and Ben-Tal, N. Conseq: the identification of functionally and structurally important residues in protein sequences. Bioinformatics, 20(8):1322–1324, 2004.

Bileschi, M. L., Belanger, D., Bryant, D. H., Sanderson, T., Carter, B., Sculley, D., DePristo, M. L., and Colwell, L. J. Using deep learning to annotate the protein universe. bioRxiv, pp. 626507, 2019.

Bitbol, A.-F., Dwyer, R. S., Colwell, L. J., and Wingreen, N. S. Inferring interaction partners from protein sequences. Proceedings of the National Academy of Sciences, 113(43):12180–12185, 2016.

Buljan, M. and Bateman, A. The evolution of protein domain families, 2009.

Burley, S. K., Berman, H. M., Kleywegt, G. J., Markley, J. L., Nakamura, H., and Velankar, S. Protein data bank (pdb): the single global macromolecular structure archive. In Protein Crystallography, pp. 627–641. Springer, 2017.

Carter, B., Mueller, J., Jain, S., and Gifford, D. What made you do this? understanding black-box decisions with sufficient input subsets. In Artificial Intelligence and Statistics, 2019.

Caruana, R., Lou, Y., Gehrke, J., Koch, P., Sturm, M., and Elhadad, N. Intelligible models for healthcare: Predicting pneumonia risk and hospital 30-day readmission. In Proceedings of the 21th ACM SIGKDD International Conference on Knowledge Discovery and Data Mining, pp. 1721–1730. ACM, 2015.

Chang, Y.-C., Hu, Z., Rachlin, J., Anton, B. P., Kasif, S., Roberts, R. J., and Steffen, M. Combrex-db: an experiment centered database of protein function: knowledge, predictions and knowledge gaps. Nucleic acids research, 44(D1):D330–D335, 2015.

Consortium, U. Uniprot: the universal protein knowledgebase. Nucleic acids research, 45(D1):D158–D169, 2016.

Dalkiran, A., Rifaioglu, A. S., Martin, M. J., Cetin-Atalay, R., Atalay, V., and Doğan, T. Ecpred: a tool for the prediction of the enzymatic functions of protein sequences based on the ec nomenclature. BMC bioinformatics, 19(1):334, 2018.

Dean, J. and Ghemawat, S. Mapreduce: simplified data processing on large clusters. Communications of the ACM, 51(1):107–113, 2008.

Deng, J., Dong, W., Socher, R., Li, L.-J., Li, K., and Fei-Fei, L. ImageNet: A Large-Scale Hierarchical Image Database. In CVPR09, 2009.

El-Gebali, S., Mistry, J., Bateman, A., Eddy, S. R., Luciani, A., Potter, S. C., Qureshi, M., Richardson, L. J., Salazar, G. A., Smart, A., et al. The pfam protein families database in 2019. Nucleic acids research, 47(D1):D427–D432, 2018.

Finn, R. D., Clements, J., and Eddy, S. R. Hmmer web server: interactive sequence similarity searching. Nucleic acids research, 39(suppl 2):W29–W37, 2011.

Golovin, D., Solnik, B., Moitra, S., Kochanski, G., Karro, J., and Sculley, D. Google vizier: A service for black-box optimization. In Proceedings of the 23rd ACM SIGKDD International Conference on Knowledge Discovery and Data Mining, pp. 1487–1495. ACM, 2017.

Gueudré, T., Baldassi, C., Zamparo, M., Weigt, M., and Pagnani, A. Simultaneous identification of specifically interacting paralogs and interprotein contacts by direct coupling analysis. Proceedings of the National Academy of Sciences, 113(43): 12186–12191, 2016.

He, K., Zhang, X., Ren, S., and Sun, J. Deep residual learning for image recognition. In Proceedings of the IEEE conference on computer vision and pattern recognition, pp. 770–778, 2016.

Hochreiter, S. and Schmidhuber, J. Long short-term memory. Neural computation, 9(8):1735–1780, 1997.

Hou, J., Adhikari, B., and Cheng, J. Deepsf: deep convolutional neural network for mapping protein sequences to folds. Bioinformatics, 34(8):1295–1303, 2017.

Kindermans, P.-J., Hooker, S., Adebayo, J., Alber, M., Schütt, T., Dähne, S., Erhan, D., and Kim, B. The (un)reliability of saliency methods. CoRR, abs/1711.00867, 2017a.

Kindermans, P.-J., Schütt, K. T., Alber, M., Müller, K.-R., Erhan, D., Kim, B., and Dähne, S. Learning how to explain neural networks: Patternnet and patternattribution. arXiv preprint arXiv:1705.05598, 2017b.

Kingma, D. P. and Ba, J. Adam: A method for stochastic optimization. International Conference on Learning Representations, 2014.

Lei, T., Barzilay, R., and Jaakkola, T. Rationalizing neural predictions. In Proceedings of the 2016 Conference on Empirical Methods in Natural Language Processing, pp. 107–117, 2016.

Li, S., Chen, J., and Liu, B. Protein remote homology detection based on bidirectional long short-term memory. BMC bioinformatics, 18(1):443, 2017a.

Li, Y., Wang, S., Umarov, R., Xie, B., Fan, M., Li, L., and Gao, X. Deepre: sequence-based enzyme ec number prediction by deep learning. Bioinformatics, 34(5):760–769, 2017b.

Liu, X. Deep recurrent neural network for protein function prediction from sequence. arXiv preprint arXiv:1701.08318, 2017.

Marks, D. S., Colwell, L. J., Sheridan, R., Hopf, T. A., Pagnani, A., Zecchina, R., and Sander, C. Protein 3d structure computed from evolutionary sequence variation. PloS one, 6(12):e28766, 2011.

McCloskey, K., Taly, A., Monti, F., Brenner, M. P., and Colwell, Using attribution to decode dataset bias in neural network models for chemistry. arXiv preprint arXiv:1811.11310, 2018.

Montavon, G., Samek, W., and Müller, K.-R. Methods for inter-preting and understanding deep neural networks. Digital Signal Processing, 73:1–15, 2018.

Price, M. N., Wetmore, K. M., Waters, R. J., Callaghan, M., Ray, J., Liu, H., Kuehl, J. V., Melnyk, R. A., Lamson, J. S., Suh, Y., et al. Mutant phenotypes for thousands of bacterial genes of unknown function. Nature, pp. 1, 2018.

R. Evans, J. Jumper, J. L. T. C. A. A. A. H. S. K. S. D. D. K. D. A. De novo structure prediction with deep-learning based scoring. Casp 13 Abstract, 2018.

Ribeiro, M. T., Singh, S., and Guestrin, C. Why should i trust you?: Explaining the predictions of any classifier. In Proceedings of the 22nd ACM SIGKDD international conference on knowledge discovery and data mining, pp. 1135–1144. ACM, 2016.

Riesselman, A. J., Ingraham, J. B., and Marks, D. S. Deep generative models of genetic variation capture the effects of mutations. Nature methods, 15(10):816–822, 2018.

Schrödinger, LLC. The PyMOL molecular graphics system, version 1.8. November 2015.

Schuster-Böckler, B., Schultz, J., and Rahmann, S. Hmm logos for visualization of protein families. BMC bioinformatics, 5(1): 7, 2004.

Seo, S., Oh, M., Park, Y., and Kim, S. Deepfam: deep learning based alignment-free method for protein family modeling and prediction. Bioinformatics, 34(13):i254–i262, 2018.

Sha, Y. and Wang, M. D. Interpretable predictions of clinical outcomes with an attention-based recurrent neural network. In Proceedings of the 8th ACM International Conference on Bioinformatics, Computational Biology, and Health Informatics, pp. 233–240. ACM, 2017.

Shrikumar, A., Greenside, P., and Kundaje, A. Learning important features through propagating activation differences. arXiv preprint arXiv:1704.02685, 2017.

Simonyan, K., Vedaldi, A., and Zisserman, A. Deep inside convolutional networks: Visualising image classification models and saliency maps. arXiv preprint arXiv:1312.6034, 2013.

Söding, J. Protein homology detection by hmm–hmm comparison. Bioinformatics, 21(7):951–960, 2004.

Springenberg, J. T., Dosovitskiy, A., Brox, T., and Riedmiller, M. Striving for simplicity: The all convolutional net. arXiv preprint arXiv:1412.6806, 2014.

Sundararajan, M., Taly, A., and Yan, Q. Axiomatic attribution for deep networks. In International Conference on Machine Learning, pp. 3319–3328, 2017.

Szalkai, B. and Grolmusz, V. Near perfect protein multi-label classification with deep neural networks. Methods, 132:50–56, 2018.

Tame, J. R. and Vallone, B. The structures of deoxy human haemoglobin and the mutant hb tyr?42his at 120 k. Acta Crystallographica Section D: Biological Crystallography, 56(7): 805–811, 2000.

Tramer, F., Zhang, F., Juels, A., Reiter, M. K., and Ristenpart, T. Stealing machine learning models via prediction apis. 2016.

Wheeler, T. J., Clements, J., and Finn, R. D. Skylign: a tool for creating informative, interactive logos representing sequence alignments and profile hidden markov models. BMC Bioinformatics, 15(1):7, 2014.

Xu, J. Distance-based protein folding powered by deep learning. arXiv preprint arXiv:1811.03481, 2018.

Yu, F. and Koltun, V. Multi-scale context aggregation by dilated convolutions. In ICLR, 2016.

Zeiler, M. D. and Fergus, R. Visualizing and understanding convolutional networks. In European conference on computer vision, pp. 818–833. Springer, 2014.

