## Supplementary Information for "Critiquing Protein Family Classification Models Using Sufficient Input Subsets"

### S1. Model Hyperparameters

We adopt the ProtCNN, 1-ResNet Block CNN, and RNN architectures from Bileschi et al. (2019) as our deep CNN, shallow CNN, and RNN models, respectively (Section 3). Our models are tuned and trained following the same procedure as Bileschi et al. (2019). Hyperparameters for each of these architectures are given in Table S1. We note that our final held-out model accuracies on the Pfam seed test set (Table 1) are similar to those presented in Bileschi et al. (2019).

*Table S1.* Hyperparameters used to train neural network models. For descriptions, see Bileschi et al. (2019).

| Hyperparameter | Deep CNN | Shallow CNN | RNN |
|---|---|---|---|
| batch size | 64 | 64 | 64 |
| dilation rate | 2 | - | - |
| filters | 450 | 500 | - |
| first dilated layer | 2 | - | - |
| gradient clip | 1 | 1 | 1 |
| kernel size | 9 | 31 | - |
| number hidden units | - | - | 1244 |
| learning rate | 0.0005 | 0.0001 | 0.0005 |
| learning rate decay rate | 0.997 | 0.997 | 0.997 |
| learning rate decay steps | 1000 | 1000 | 1000 |
| learning rate warmup steps | 3000 | 3000 | 3000 |
| number of ResNet layers | 5 | 1 | - |
| pooling | max | max | mean |
| ResNet bottleneck factor | 0.5 | 0.5 | - |
| train steps | 400000 | 400000 | 300000 |
